# Basal Cell-Contact Dynamics Influence Tissue Packing in a Proliferating Mammalian Epithelium

**DOI:** 10.1101/2025.07.30.665610

**Authors:** Subramanian P. Ramanathan, Tanmoy Sarkar, Matej Krajnc, Rosemary Mwithiga, Azize Cerci, Chongbei Zhao, Matthew C. Gibson

## Abstract

Animal tissue morphology is determined by the shape, position, and proliferative capacity of individual epithelial cells. Nevertheless, it remains incompletely understood how the dynamic shape transformations implicit in mitotic proliferation influence tissue packing, particularly at the level of basal cell contacts. Here, we use an *in silico* vertex model to show that epithelial mitotic rounding necessitates a sequence of dynamic basal contact rearrangements, including basal diminution of the mitotic cell volume, transient multicellular rosette assembly, basal reinsertion of daughter cells, and neighbor reorganization. We then leverage a mammalian intestinal organoid model to confirm nearly identical basal cell-contact dynamics as those predicted *in silico*. Pharmacological inhibition of mitotic progression reveals that two events—basal diminution of the cell body and daughter cell reinsertion—independently drive distinct contact rearrangements. Together, our results uncover a previously underappreciated topological role for basal mitotic cell dynamics in shaping epithelial packing and morphogenesis.

## INTRODUCTION

Organogenesis requires precise control of cell packing and cell numbers, which is governed by cell position and proliferation, respectively. Being tightly organized and otherwise mostly immotile, epithelial cells depend on distinct cell contact transformations and consequent neighbor exchanges to reposition individual cells (Bertet et al., 2004; Classen et al., 2005; Fernandez-Gonzalez et al., 2009). Distinct modes of epithelial cell reorganization are driven by characteristic contact transformations. For instance, neighbor cell exchanges are achieved by contracting existing cell contacts and extending new contacts (T1 and rosettes), whereas cell shrinkage and elimination is accommodated by simultaneously contracting all of the cell’s edges (T2) (Bertet et al., 2004; Blankenship et al., 2006; Guillot and Lecuit, 2013). New cells can be incorporated either by establishing *de novo* contacts between daughter cells after division or by forming multiple new contacts during radial migration events such as apical emergence (Gibson et al., 2006; Sedzinski et al., 2016; Christodoulou and Skourides, 2025). Most models of cell contact transformation dynamics and cell repositioning have primarily been limited to analyzing the differential tension and adhesion in apical adherens junctions, assuming that the apical cell-contact architecture will be recapitulated basally (Lemke and Nelson, 2021). However, more recent work shows that cell contact topology and dynamics are strongly influenced by cell and tissue geometry Rupprecht et al., 2017; Gómez-Gálvez et al., 2018). Consequently, differences in cell morphology between the apical and basal sides can give rise to distinct cell contact behaviors at each surface (Rozman et al., 2024). Nevertheless, an understanding of basal cell contacts and cell positioning is still limited, especially in proliferating tissues.

Mitotic cells adopt an asymmetric position along the apico-basal axis of epithelia. At mitotic entry, cells undergo a dramatic morphological transformation whereby their cellular mass translocates apically and the cell assumes a spherical shape (Sauer, 1935). This transformation leads to a near-complete loss of basal volume, a phenomenon we refer to as basal diminution. This mitotic shape change is evolutionarily conserved across proliferating epithelia (Meyer et al., 2011) and influences both development and disease in diverse tissue types (Fleming et al., 2007; Luxenburg et al., 2011; Nakajima et al., 2013; Kondo and Hayashi, 2013; Rosa et al., 2015; Hoijman et al., 2015; Freddo et al., 2016; Chanet et al., 2017; Petridou et al., 2019; Aguilar-Aragon et al., 2020; Despin-Guitard et al., 2024). Furthermore, it has been shown that the geometry of nascent apical junctions between daughter cells directly influences epithelial cell packing to ensure a consistent cell shape distribution within tissues (Gibson et al., 2006, 2011) and can also influence cell rearrangement (Firmino et al., 2016; Petridou et al., 2019; Godard et al., 2020). Together, these observations highlight the well-established role of mitotic cell shape and cell-contact topology in development and disease, particularly at the apical surface. However, the impact of mitotic cell shape dynamics on basal cell contact topology is largely unknown.

## RESULTS

### Vertex model simulations of epithelial dynamics predict basal neighbor exchanges during mitosis

The mechanical behaviors associated with mitotic rounding and changes in apical contact topology have been well characterized using direct force measurements and computational modeling (Gibson et al., 2006, 2011; Ragkousi and Gibson, 2014; Ramanathan et al., 2015, 2019). However, our understanding of the mitosis-associated morphological and topological dynamics in the basal plane of epithelial tissues is limited. To address this, we employed 2D vertex model simulations (Farhadifar et al., 2007; Alt et al., 2017; Sugimura and Otani, 2024) to examine basal contact dynamics during cell division. To mimic apical mitotic rounding (Meyer et al., 2011), simulated mitotic cells were progressively removed from the basal plane and subsequently reinserted as two co-equal daughter cells (Figure 1A, B). Assuming the average cell size in the tissue is approximately 50 µm2, we set the value of the preferred perimeter (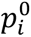) in the vertex model to 27.22 µm, which corresponds to a shape factor (perimeter/area^1/2^) of ∼ 3.85 (Figure 1). Strikingly, the removal and reinsertion of mitotic cells frequently triggered basal neighbor exchanges and cell mixing (Figure 1C). Notably, over 88% of T1 transitions (neighbor exchanges) occurred during daughter cell reinsertion, rather than during the initial basal diminution phase (Figure 1D). Although mitosis-driven T1s were observed throughout the epithelium, their spatial distribution within the tissue was heavily skewed towards cell contacts closer to the mitotic cell (Figure 1E). This trend held across a range of preferred perimeters representing tissue regimes that were either solid-like (e.g., 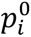 = 26.16 µm, shape factor ∼ 3.81) or floppy (e.g., 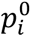 = 28.28 µm, shape factor ∼ 4.00; and 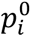 = 29.34 µm, shape factor ∼ 4.15; Figure S1A). In all cases, T1 transitions predominantly occurred shortly after and in close proximity to daughter cell reinsertion (Figure S1B, C). Unlike previous *in silico* models that simulate mitotic division by bifurcating a single mother cell, our approach captures the dynamics of basal topology, predicting sequential events: basal cell diminution/elimination (T2-like transitions), daughter cell reinsertion, and mitosis-driven neighbor exchanges (T1s). These results predict a clear contrast between apical and basal cell contact dynamics during epithelial cell division, which prompted us to validate the model in living tissue.

**Figure 1.**
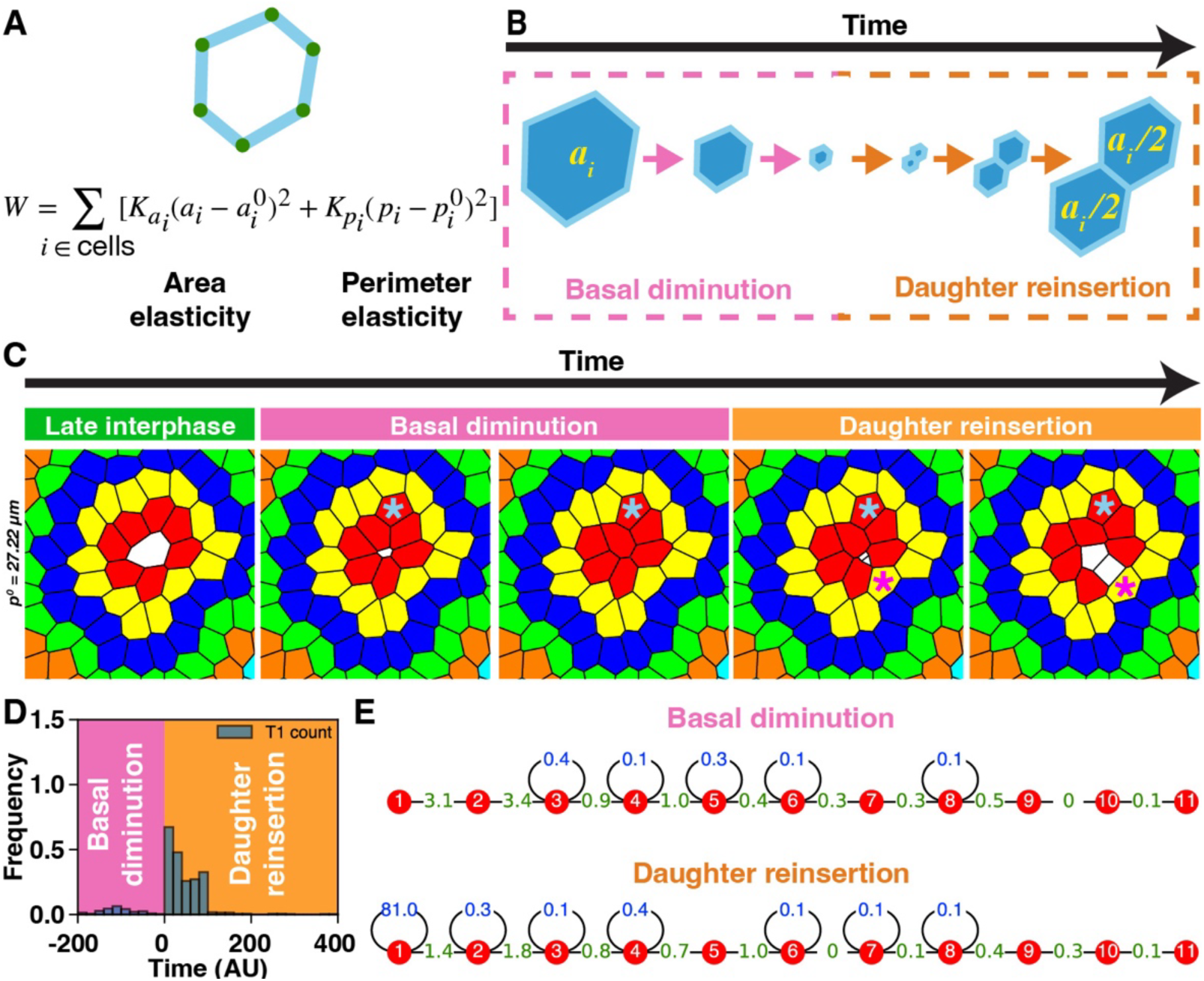
Computational simulations of mitosis predict mitosis-associated basal contact remodeling and neighbor exchanges. (A) Schematic of the *in silico* vertex model in which epithelial cells are represented as polygons defined by vertices (green) and straight edges (cyan). Tissue mechanics are governed by an energy function, *W*, consisting of area elasticity (with elastic constant *K*_*a*_) and perimeter elasticity (with constant *K*_*p*_), each with corresponding preferred values 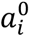 and 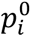. Cells with larger preferred perimeters adopt more elongated shapes. Vertex positions evolve to minimize *W*. (B) Cell division is simulated in two phases: (1) basal diminution — modeled as a gradual reduction in the preferred area and perimeter of the mitotic cell, effectively removing it from the basal plane; and (2) reinsertion — in which two daughter cells are reinserted with half the preferred area of the mother cell. (C) Snapshots showing basal contact rearrangements during mitosis. The mitotic cell (0°) is white, and the neighboring cells (1°-6°) are color-coded by topological proximity. The preferred perimeter (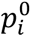) of interphase cells are set to 27.22 µ*m*. Cyan and magenta asterisks denote neighbor exchanges during mitotic cell diminution and daughter cell reinsertion, respectively. (D) Temporal distribution of T1 transitions (*n* = 324 divisions), with negative and positive times corresponding to diminution and reinsertion, respectively. T1s on the edges of the mitotic and the daughter cells are excluded. (E) Spatial distribution of the T1 transitions as a function of topological distance (red nodes) from the mitotic cell. The numbers indicate the percentage of total T1 transitions occurring throughout the entire cell division process. T1s between cells at the same or different topological distances are shown in blue and green, respectively.

### Mitotic cell morphology and dynamics in intestinal organoid epithelium

In most animal tissues, the basal surfaces of polarized epithelia are tightly associated with apposing cell layers or a thickened basement membrane, making high-resolution imaging of basal cell dynamics technically challenging. To overcome this, we leveraged mouse intestinal organoids, wherein the apical epithelial plane forms at the interior of the tissue, enclosing a central lumen (anti-ZO-1; Figure S2A). Consequently, the basal plane is exposed to the exterior and accessible for imaging studies (anti-Laminin; Figure S2B). We first confirmed that the hallmark mitotic cell behaviors observed *in vivo* are preserved in this model. Specifically, key features such as the apical displacement and rounding of mitotic cell bodies and the subsequent reinsertion of daughter cells into the basal region of the epithelium were clearly recapitulated (Figure S2C). Both fixed and live imaging demonstrated that organoid cells form a columnar epithelium with interphase nuclei localized near the basal surface (Figure S2C, 2A). As observed in intact mammalian intestine, upon breakdown of the nuclear envelope at the start of mitosis, the cellular mass translocated away from the basal surface during mitotic rounding (Trier, 1963; Fleming et al., 2007). After completing mitosis apically, daughter cells then reinserted into the basal region (Figure 2A). Together, the conservation of epithelial cell dynamics and the accessibility of the exposed basal plane clearly establish intestinal organoids as an ideal model for tracking basal topology during mitosis.

**Figure 2.**
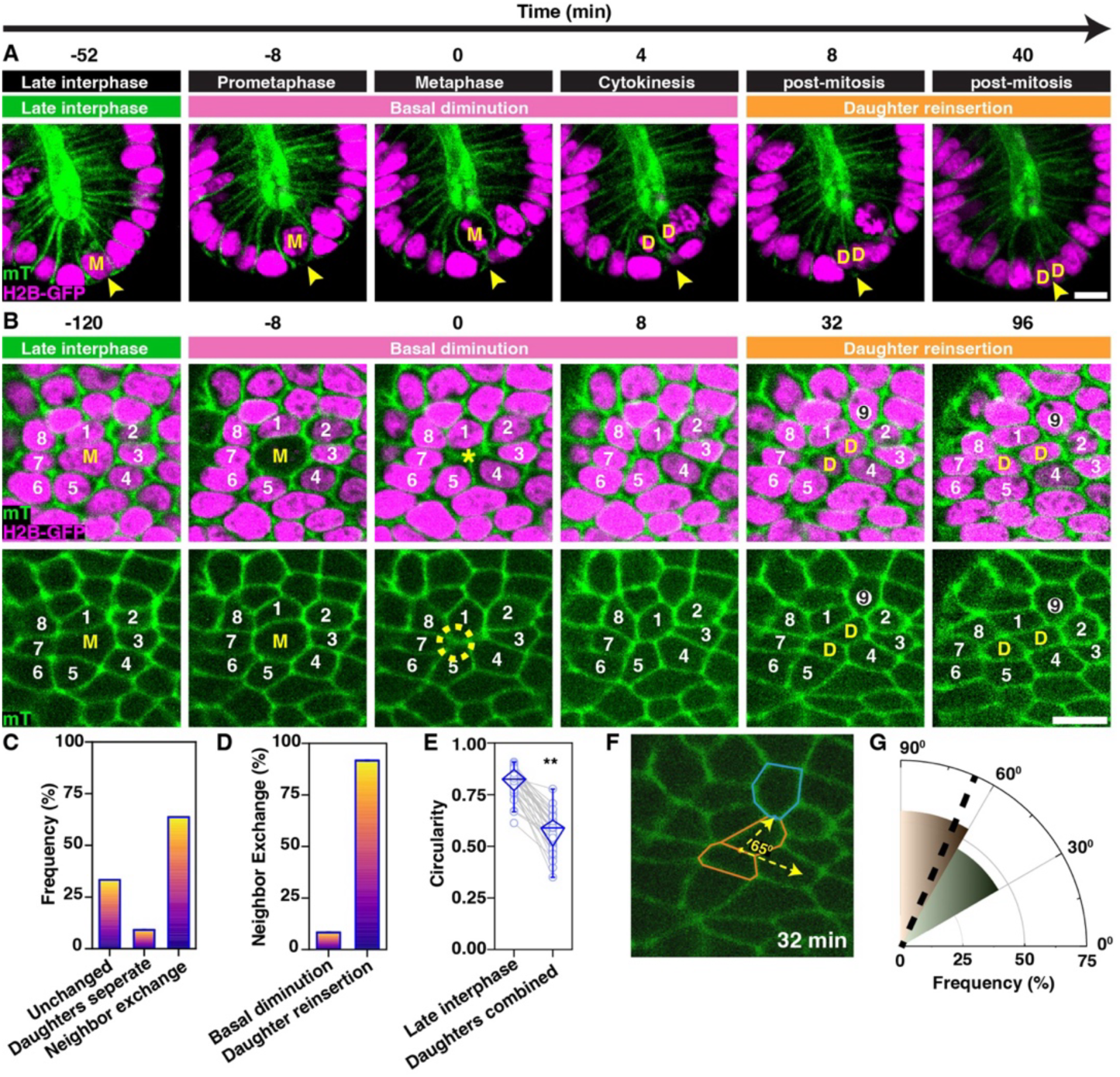
Basal diminution of the mitotic cell and the reinsertion of daughter cell body coincide with local cell shape changes, basal rosettes, and neighbor exchanges. (A) Timelapse images of the intestinal organoid epithelium showing the morphological changes of a mitotic cell (**‘M’**) and daughter cells (**‘D’**), including basal diminution, mitotic cell rounding, cytokinesis, and basal reinsertion. The sites of basal diminution and reinsertion are indicated by arrowheads. Cell membranes: mTomato (green); nuclei: H2B-GFP (magenta). Scale bar, 10 µm. (B) Cross-sectional images showing the basal cell-contact topology of a mitotic cell (‘**M’**), its immediate neighbors (‘**1-8’**), and reinserting daughter cells (‘**D’**). The interposition of a new immediate neighbor (cell ‘**9**’) separated cells ‘**1**’ and ‘**2**’. The yellow circle highlights a transient rosette with five cells sharing a vertex. Scale bar, 10 µm. (C) Frequency of daughter cell separation versus to local neighbor exchange (*n* = 33 mitoses). (D) Proportion of mitosis-associated 1^0^ neighbor exchanges during diminution vs. daughter reinsertion. (E) Circularity of mother cell in late interphase vs. combined daughters post-division. ** denotes *p*-value < 0.05, Paired Wilcoxon Signed Ranks Test. Unless otherwise specified, in this and every following box plot, circles represent individual cells, the diamond box contains 25–75% percentiles of the data, and the bar denotes the median. (F) Schematic measuring the angle between the cell contact shared between the daughter cells (orange) and that of the new neighbor (teal). (G) In the polar graph, the petals indicate the frequency of the angle between the cell contact shared between the daughter cell and that of the new neighbor. Median indicated by a radial line.

### Basal cell-contact transformations during mitosis in organoid epithelia

To investigate the topological dynamics at the basal surface during epithelial mitosis, we imaged dividing cells in mouse intestinal organoids. At mitotic entry, the basal cell-contact topology resembled a T2-like event, characterized by the progressive diminution and eventual disappearance of the mitotic cell from the basal plane (Figure 2B). This was frequently accompanied by the formation of a transient multicellular rosette-like contact configuration. In most cases (∼73% of 33 cell divisions), the subsequent reinsertion of daughter cells was accompanied by basal neighbor exchanges and local cell mixing. Consistent with previous reports (Carroll et al., 2017; McKinley et al., 2018), daughter cells occasionally separated during reinsertion (∼9% of events; Figure S3, Figure 2C). However, the majority of daughter cell reinsertion events were associated with contact exchanges between cells in the immediate neighborhood (64%; Figure 2B-D). While the combined basal area of daughter cells was comparable to that of their mitotic mother in late interphase (Figure S4), we did observe a consistent increase in the basal perimeter of reinserted daughter cells compared to that measured in the corresponding late interphase cells. Indeed, the circularity of the two daughters combined was 30% less than that of the progenitor in late interphase, indicating that the shape of the two daughter cells combined was more elongated (Figure 2E). Interestingly, changes in contact topology and neighbor exchanges were more likely to occur perpendicular to the shared junction between the daughter cells (Figure 2F, G). Together, these observations reveal that at the basal plane, mitosis involves a sequence of morphological and topological changes: cell body diminution (T2-like transitions), apical division, basal reinsertion of daughter cells, and mitosis-driven neighbor exchanges (T1 transitions). These findings closely align with predictions from our vertex model simulations (Figure 1). However, while rosette-like contact patterns were a consistent feature of *in vivo* divisions, they were not observed *in silico*.

### The incomplete elimination of mitotic cells from the basal plane generates rosette-like cell contacts *in silico*

Multicellular rosettes are widely observed apical structures that bring into contact four or more epithelial cells at a single vertex, typically by elevating contractility at the level of the adherens junctions (Blankenship et al., 2006). These configurations are inherently unstable and serve as transient intermediate topological arrangements in a range of organs and organisms (Harding et al., 2014; Neahring and Zallen, 2025). Intriguingly, we observed rosette-like cell assemblages at the basal regions of proliferating organoid epithelia (Figure 2B). In diverse tissues, including the mammalian intestine, apically displaced mitotic cells often retain attachment to the basement membrane through thin, elongated cellular tethers typically less than 100 nm in diameter (Sauer, 1935; Fleming et al., 2007; Kosodo et al., 2008; Carroll et al., 2017).

To explore the potential contribution of mitotic cell tethers in stabilizing basal rosettes and inducing neighbor exchanges, we used vertex model simulations. Specifically, we modeled a mitotic cell that was incompletely eliminated from the basal plane, leaving behind a tether either one-millionth or one-hundredth of the initial size of the interphase cell (Figure 3A, B). When mitotic cell remnants were vanishingly small (one-millionth the original area), basal contact dynamics were indistinguishable from those observed with complete elimination. However, when larger remnants (∼1% of interphase cell area) were retained, rosette-like arrangements of neighboring cells consistently formed (Figure 3B). Notably, the presence of these rosettes did not lead to an increase in neighbor exchanges (Figure 3B, C). On the contrary, simulations across a range of mechanical regimes—spanning solid-like to floppy tissues (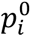 = 26.16 µm, 27.22 µm, 28.28 µm, and 29.34 µm)—consistently showed a reduction in T1 transitions during daughter cell reinsertion when rosettes were present (Figure 3D). These results suggest that basal rosettes can arise passively from persistent mitotic tethers but that their formation suppresses, rather than promotes, neighbor exchanges. Thus, while structurally similar to apical rosettes, basal rosettes formed through incomplete cell elimination are unlikely to serve as active drivers of epithelial remodeling.

**Figure 3.**
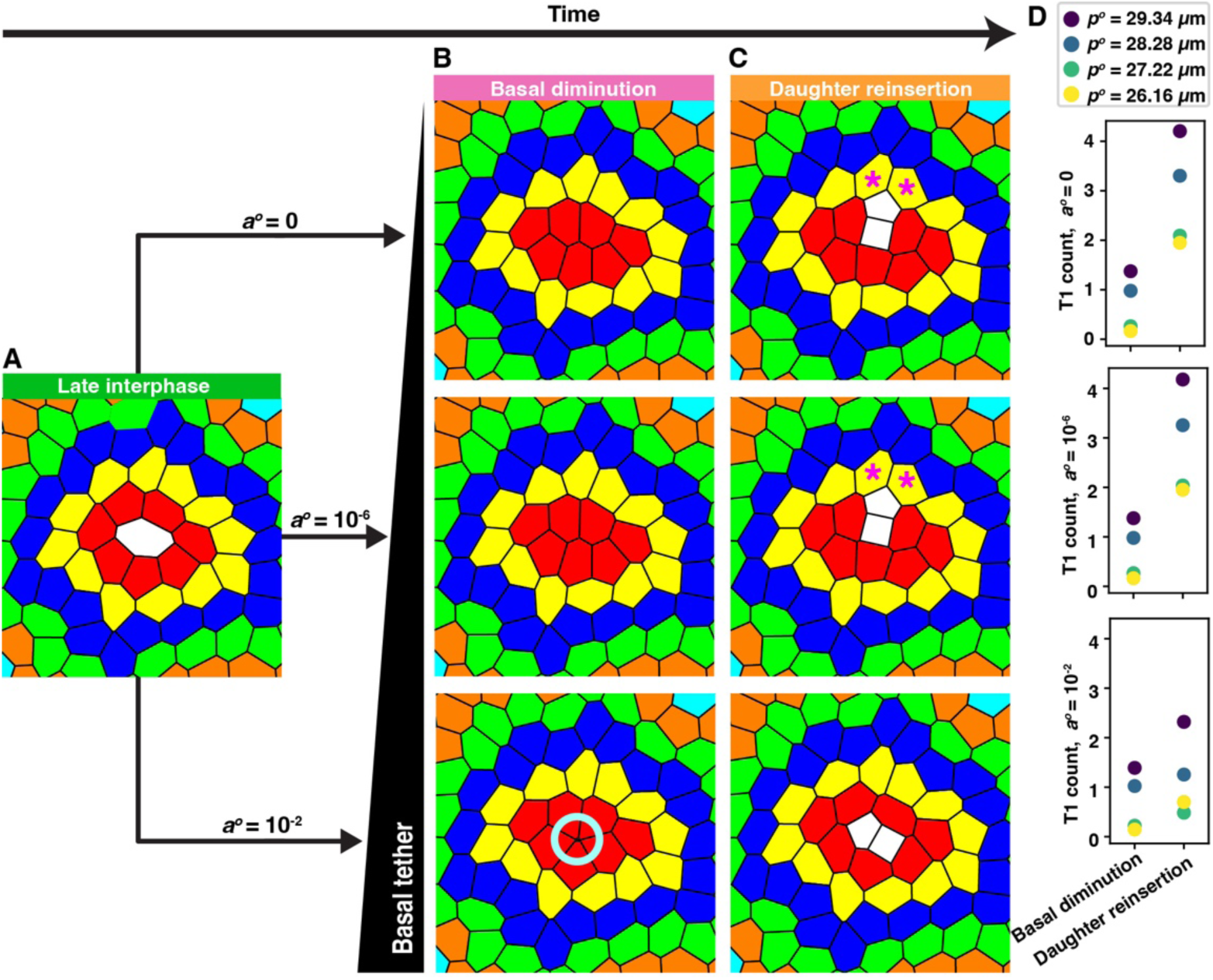
Simulated mitotic tethers generate basal rosettes but suppress contact rearrangements. (A) Snapshot of a dividing cell (white) in a tissue with 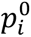 = 27.22 µ*m*. (B) Mitotic cells are completely (*a*^0^ = 0µ*m*^2^) or partially (*a*^0^ = 5*10^−5^*μm*^2^and 5*10^−1^*μm*^2^) displaced from the basal plane. Incomplete diminution with *a*^0^ = 5*10^−1^*μm*^2^ produced rosette-like structures (teal circle). (C) Post-division reinsertion of daughter cells. Magenta asterisks mark examples of T1 transitions. (D) T1 transition frequency under each diminution condition across tissues of varying 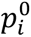 (26.16*μm* - 29.34*μm*). n = 324 divisions per condition.

### The deformation of local tissue is a consequence of daughter cell reinsertion

Having established that basal rosette-like structures are unlikely to contribute to neighbor exchanges during mitosis, we next examined the role of daughter cell insertion in influencing cell contact topology. Both *in silico* simulations and live imaging of organoid epithelia revealed that the majority of neighbor exchanges were temporally coupled with the reinsertion of post-mitotic daughter cells into the basal plane (Figure 1D; Figure 2D). Moreover, these exchanges were predominantly oriented orthogonally to the newly formed cell contact between daughter cells (Figure 2F, G). To test whether the orientation of reinsertion influences the directionality of neighbor exchanges, we analyzed vertex model simulations across tissues with varying mechanical properties—spanning both solid-like and floppy regimes (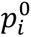 = 26.16 µm, 27.22 µm, 28.28 µm, and 29.34 µm). Across all simulations, fewer than 8% of neighbor exchanges occurred within 30° of the axis aligned with the nascent daughter cell contact, confirming a strong bias for orthogonal reorganization (Figure 4A, B). Together, these data demonstrate a robust spatiotemporal correlation between basal neighbor exchanges and the reinsertion of daughter cells following mitosis.

**Figure 4.**
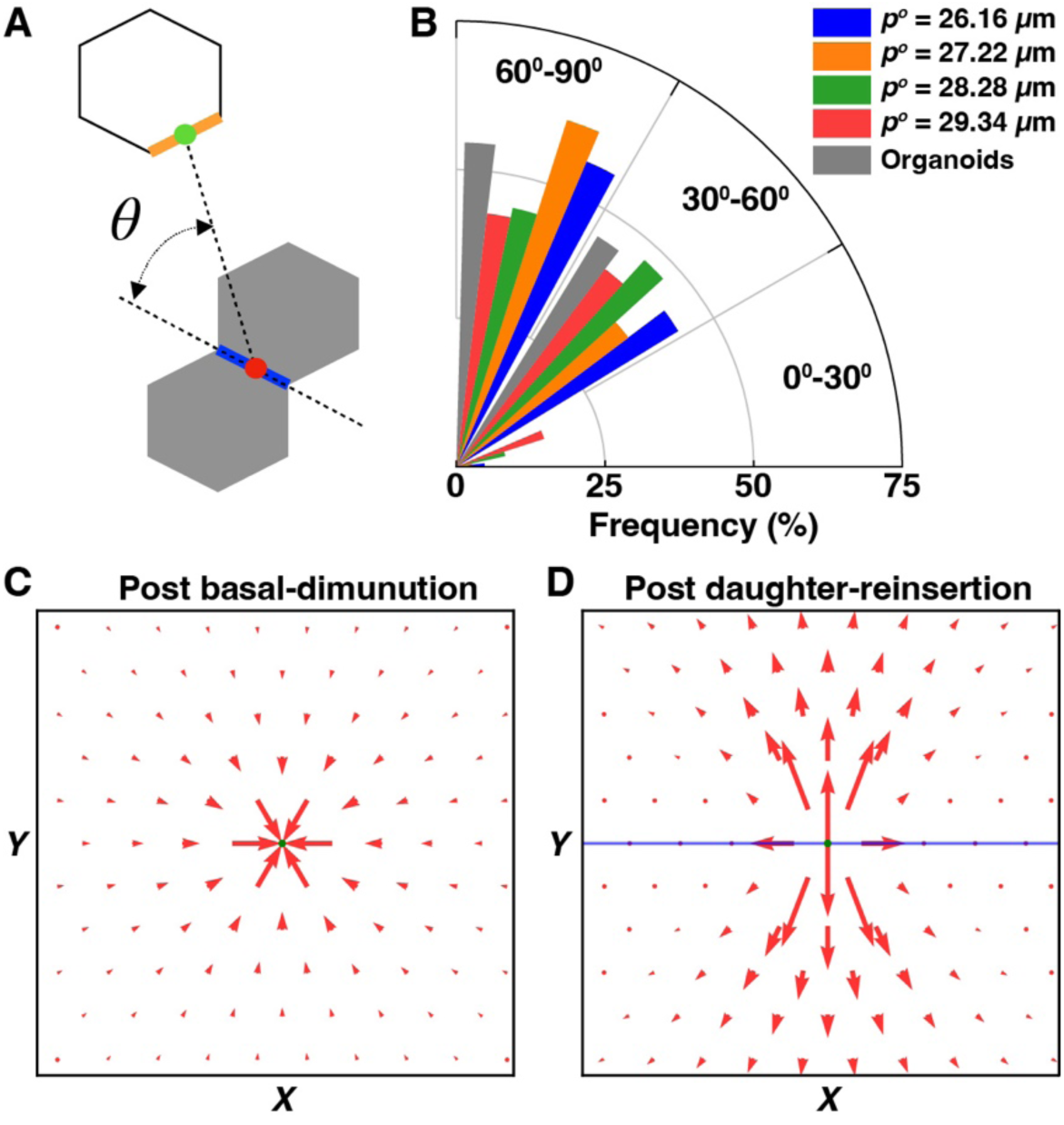
The reinsertion of daughter cells generates anisotropic strain and drives neighbor exchange orientation. (A) Schematic measuring angle between the daughter-daughter contact (blue) and the newly formed T1 cell contact (orange). (B) Polar histogram showing the orientation of T1 cell contacts *in silico* (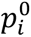= 26.16*μm* - 29.34*μm*) and in organoids (grey). *n* = 324 and 22 divisions *in silico* and *in vivo*, respectively. (C, D) Displacement vector fields of cell centroids during (C) diminution and (D) reinsertion. Arrow length and direction indicate magnitude and orientation of local displacement. The blue segment is colinear with daughter cell contacts.

To investigate the mechanical basis of this correlation, we quantified local tissue strain during the mitotic cycle. Starting from a defect-free honeycomb lattice, we tracked the displacement of cell centroids during basal diminution and subsequent reinsertion. During diminution, displacements were greatest in the immediate vicinity of the mitotic cell but remained isotropic in orientation (Figure 4C). In contrast, reinsertion of daughter cells generated anisotropic displacements, oriented perpendicular to the cell contact shared by the daughters (Figure 4D). Together, these results suggest that anisotropic local strain, generated during daughter cell reinsertion, is the primary mechanical driver of mitosis-associated neighbor exchanges at the basal surface.

### Neighbor exchanges in the basal region of proliferating organoid epithelium require the basal reinsertion of daughter cells

Thus far, our results closely follow the predictions *in silico* and show that reinsertion of daughter cells spatially and temporally correlates with T1 transitions *in vivo* (Figure 2B-G). We next sought to experimentally test whether daughter cell reinsertion is essential for driving neighbor exchanges by preventing mitotic cells from reinserting into the basal plane. S-trityl-l-cysteine (STC) treatment can arrest cells in mitosis without altering their mechanical properties (Skoufias et al., 2006; Ramanathan et al., 2015). Treating intestinal organoids with STC caused mitotic cell bodies to be held at the apical region and thus indefinitely depleted from the basal plane (Figure 5A-C). In addition to maintaining their apical localization, mitotic cells also maintained their spherical shape for durations of >1h (Figure 5B). Furthermore, the morphology of late interphase cells and their neighbors did not change with STC treatment, further indicating that the general mechanical properties of cells were unperturbed (Figure 5D, E). This is consistent with prior mechanical measurements showing that STC arrests cells in mitosis without altering their intrinsic stiffness or shape (Ramanathan et al., 2015). Changes in the mechanical and morphological properties of cells have been shown to change epithelial packing by altering cell-neighbor relationships (Bardet et al., 2013; Ramanathan et al., 2019). In close agreement with observations from several epithelial tissue types (Gibson et al., 2006), late interphase cells that entered mitosis within an hour and their neighboring cells had a median number of neighbors of seven and six, respectively (Figure 5F). The fact that STC treatment did not change the expected distribution of epithelial neighbors indicates that the mechanical property of organoid tissue was not altered.

**Figure 5.**
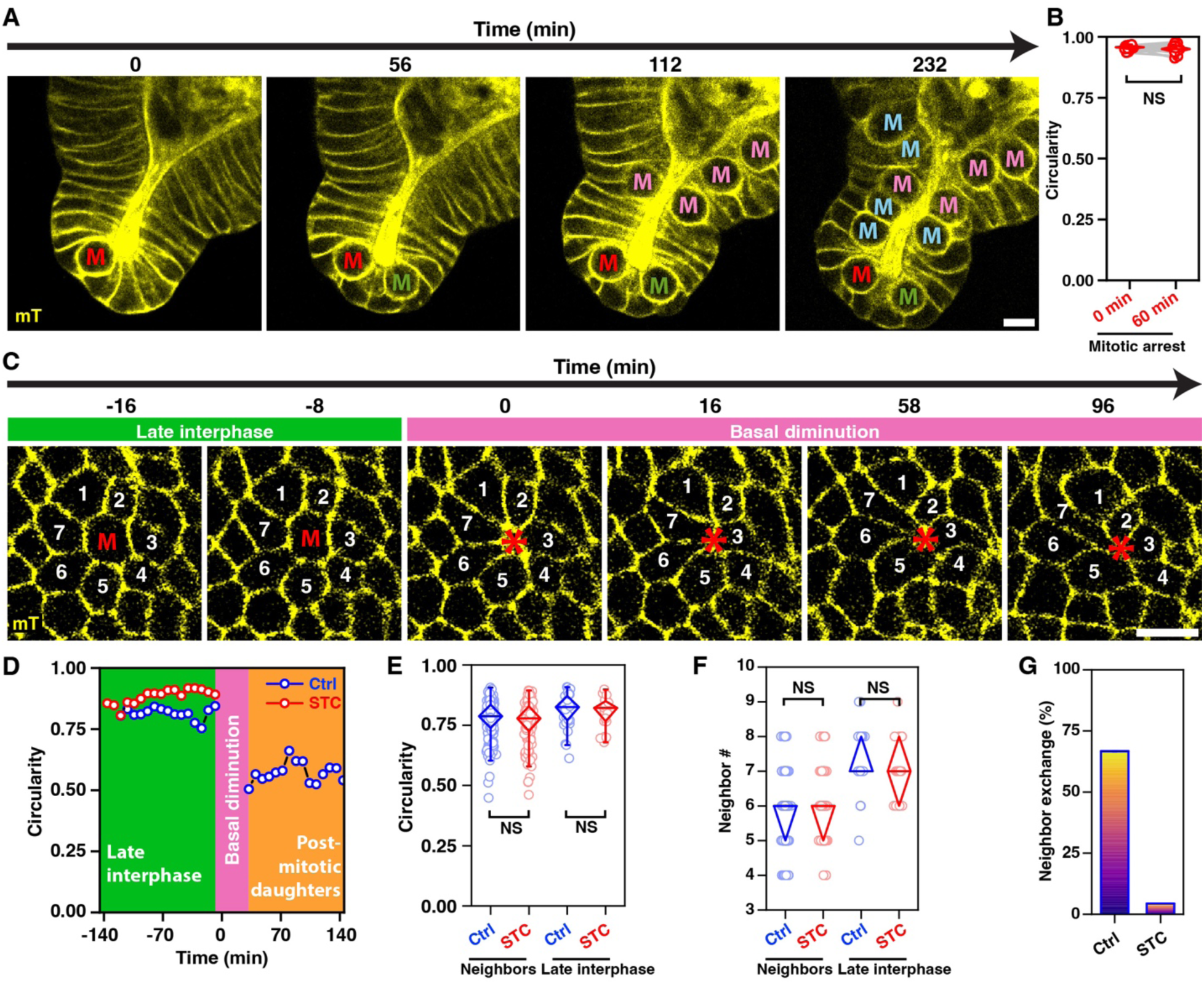
Abolishing the reinsertion of daughter cells blocks neighbor exchanges. (A) Timelapse images of an STC-treated organoid showing the morphological changes associated with cells entering and remaining in mitosis (‘**M’**). Red, green, magenta, and cyan lettering denote mitotic entry at 0, 56, 112, and 232 min, respectively. Cell membrane: mTomato (yellow). All scale bars, 10 µm. (B) Circularity of apically located spherical mitotic cell 0 min and 60 min after rounding (*n* = 15 cells). NS denotes *p*-value > 0.05, Wilcoxon Signed Ranks Test. (C) Cross-sectional images showing the basal cell-contact topology of an arrested mitotic cell (‘**M’**) and its immediate neighbors (‘**1-7’**). (D) Circularity of untreated (blue) vs. STC-treated (red) cells before mitosis (green background), during basal diminution (pink background), and after daughter reinsertion (orange background). The data for STC-treated mitotic cell are limited to late interphase because they do not reinsert into the basal plane after basal diminution. (E, F) Cell circularity (D) and number of neighbors (E) of untreated (blue) and STC-treated (red cells) pre-mitotic cells and their immediate neighbors. *n* = 31 untreated and 23 STC-treated mitotic cells. NS denotes *p*-value > 0.05, Mann-Whitney Test. (G) Frequency of local neighbor exchanges in untreated and STC-treated organoids.

Having validated that using STC treatment to arrest apically rounded cells in mitosis does not alter their mechanical and morphological properties, we next investigated the topological consequence of blocking daughter cell reinsertion. Consistent with observations from untreated controls, STC-treated organoids exhibited transient basal-rosette-like structures associated with cells undergoing apical rounding (Figure 5C). Nevertheless, compared with untreated controls, cell exchanges among the first-order neighbors were reduced by 93% in STC-treated organoids (Figure 5C, G). These results demonstrate that daughter cell reinsertion is required for cell-neighbor exchanges in proliferating intestinal epithelia. As cell shapes and the mechanical properties of the epithelium were unaltered by STC treatment, we conclude that the basal neighbor exchanges are mechanically driven by daughter cell reinsertion. Altogether, these *in vivo* results are in close agreement with our *in silico* findings, demonstrating that a basal T2-like cell elimination process and the formation of basal rosette-like cell contacts are associated with basal diminution, while daughter cell reinsertion drives local rearrangements and neighbor exchanges (T1 transitions) at the basal plane.

## DISCUSSION

Tissue geometry and mechanics are well known to influence mitotic progression and daughter cell dynamics in epithelia. For example, the local topology of neighboring cells can orient the mitotic cleavage plane across a wide variety of systems, including plants and animals (Gibson et al., 2011). Similarly, local mechanical constraints can determine whether daughter cells remain adjacent or are separated during reinsertion, as demonstrated in the chick epiblast (Firmino et al., 2016) and mammalian intestinal organoids (Carroll et al., 2017; McKinley et al., 2018). Our findings demonstrate that the converse is also true: mitotic cell shape changes can drive topological remodeling of the surrounding epithelium. At the basal surface, mitotic cells not only undergo a bifurcation to produce daughter cells but also trigger a sequence of transformations in their neighbors—including T2-like diminution, rosette formation, and T1 neighbor exchanges. Notably, most neighbor exchanges occur not during the early phase of mitotic cell disappearance from the basal plane, but during the reinsertion of daughter cells. This transition is marked by the emergence of non-isometric, anisotropic daughter cell shapes that appear to locally deform the tissue, generating biased strain patterns. These mechanical asymmetries – apical displacement orthogonal to and daughter insertion within the epithelial plane – are sufficient to orient local packing transformations. Taken together, these findings emphasize that mitosis is not a topologically passive process but a mechanically active event that sculpts epithelial topology.

Basal diminution during mitosis raises interesting parallels with another key driver of epithelial remodeling: apoptosis. Classically, apoptosis is modeled as a T2 transition involving isotropic cell-contact contraction and neighbor remodeling (Alt et al., 2017). However, recent studies suggest that epithelial apoptosis shares several mechanistic features with mitosis. For instance, apoptotic cells exhibit cytoskeleton-driven, asymmetric repositioning of the nucleus and cell body along the apico-basal axis, as well as the persistence of nanoscopic cellular remnants embedded within the epithelial layer after cell body loss (Arnould et al., 2025). Like mitosis, the apoptosis can influence local tissue organization, driving epithelial closure (Toyama et al., 2008) or folding and bending (Monier et al., 2015; Ambrosini et al., 2019). These shared features suggest that, just as daughter cell reinsertion actively drives neighbor exchanges during mitosis, apoptotic extrusion may also trigger remodeling shaped by local geometry, strain anisotropy, and packing constraints. Future studies using vertex modeling and live imaging— incorporating factors like tether persistence, anisotropic strain, and cell contact remodeling—could reveal that apoptosis, like mitosis, plays an active and instructive role in shaping epithelial topology.

We find that rosette formation—a hallmark of apical remodeling—can also emerge transiently at the basal surface during mitosis, representing a spatially distinct and previously underappreciated intermediate in epithelial reorganization. Notably, in *Drosophila*, complex apical junctional structures such as rosettes have been shown to initiate from basal contacts (Sun et al., 2017), and similar multicellular rosettes play central roles in organizing tissue morphogenesis across species. Although the precise initiation mechanisms remain unclear, mitotic cell shape changes are increasingly implicated in triggering rosette formation—for example, in the left-right organizer of zebrafish and mice (Abdel-Razek et al., 2023; Wu et al., 2025). Furthermore, in the mouse limb bud, anisotropic stress aligns cell divisions and rosettes to remodel tissue geometry (Lau et al., 2015). While apical rosettes have been extensively studied, our findings suggest that basal rosettes may represent a distinct and functionally relevant motif during mitosis. However, computational simulations indicate that when basal rosettes are stabilized by persistent tethers or incomplete cell elimination, they do not facilitate neighbor exchanges and may even delay them. These observations suggest that although rosettes are a conserved morphological feature, their functional impact likely depends on spatial context, mechanical constraints, and cell cycle stage.

Looking forward, many fundamental questions remain. How do local tissue mechanics—such as tension anisotropy, planar polarity, and curvature—influence basal mitotic behaviors? How does basement membrane composition or stiffness affect reinsertion dynamics and tissue deformation? Do different epithelial subtypes or developmental stages exhibit unique strategies for coordinating mitosis with tissue remodeling? Addressing these questions will require the integration of in vivo imaging, quantitative modeling, and mechanobiological perturbations. In conclusion, our work reveals that mitosis is a potent driver of topological transformation at the basal surface of epithelia. The interplay between cell shape, mechanical asymmetry, and neighbor connectivity suggests a broader framework where proliferation is not merely additive but morphogenetic—continuously shaping and refining epithelial architecture. Revisiting other cellular transitions, including apoptosis, with this mechanical and topological lens may further enrich our understanding of how tissues self-organize during development, regeneration, and disease.

## MATERIALS AND METHODS

### Lead Contact and Materials Availability

Further information and requests for resources and reagents should be directed to and will be fulfilled by the Lead Contact, Subramanian P. Ramanathan.

### Experimental Models and Subject Details

#### *In silico* simulations

##### Vertex model

In two-dimensional (2D) vertex models, tissues are represented as planar tilings of polygonal cells. The shape of a polygonal cell is parametrized by the location of its vertices **r_v_** = (*x*_*v*_, *y*_*v*_), where *v* stands for the *v*-th vertex of the system. Any cellular scale dynamics, such as cellular deformations or displacements, are captured through the motion of these vertices governed by the overdamped dynamics:

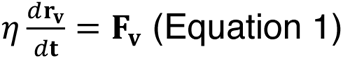

where *η* is the friction coefficient of vertices against the substrate, and *F*_*v*_ is the net force acting on the vertex *v*. The force on a vertex is derived from a conservative potential energy that always draws the system towards its local minima. Here, the potential energy contains contributions from area and perimeter elasticity as follows

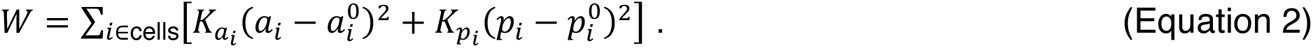

Here, *a*_*i*_ and *p*_*i*_ are the actual area and perimeter of the *i*-th cell, 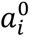 and 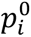 are the corresponding preferred values, and 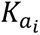 and 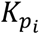 are the moduli associated with area and perimeter elasticity, respectively. The force on each vertex is given by ***F***_*v*_ = −∇_*v*_W, where 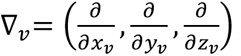. We consider that *a*_0_ is the area of any normal cell, and 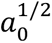 is the unit of length. Additionally, we assume identical elastic properties for all cells, i.e., 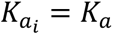 and 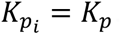.

We nondimensionalize the system by defining dimensionless variables: 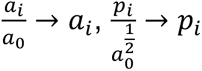, 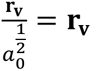, and 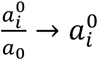. The resulting dimensionless equation of motion for the *v*-th vertex reads

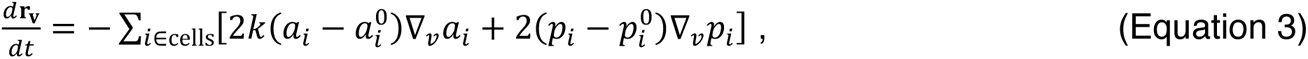

where 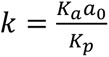 is the dimensionless area elasticity modulus, whereas 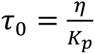 is the characteristic time. Furthermore, we assume that the cells are incompressible and set *k* = 100. Using a forward finite-difference scheme, we use a time step Δ*t* = 10^−3^ to solve the Eq. (3).

##### Initial configuration preparation

We take a pre-generated disordered 2D tissue with *N*_*c*_ = 324 equal-sized cells 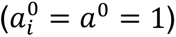 under periodic boundary conditions. From this base configuration, we generate *N*_*c*_ configurations by designating one arbitrarily chosen cell from each as a large mitotic cell with preferred area 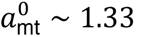. To accommodate this increase, we adjust the system size accordingly.

Additionally, we choose four values for the preferred perimeters of normal cells: 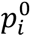 = *p*^0^ = 3.70, 3.85, 4.00, and 4.15, resulting in four distinct sets of configurations, each containing *N*_*c*_ samples. These dimensionless values correspond to physical preferred cell areas (*a*^0^) of approximately 50 *μm*^2^, and perimeters of 26.16*μm*, 27.22*μm*, 28.28*μm*, and 29.34*μm* corresponding to *p*^0^ = 3.70, 3.85, 4.00, and 4.15, respectively.

Each configuration is then melted using an active-tension-fluctuations scheme, described by the Ornstein-Uhlenbeck model (Curran et al., 2017; Krajnc, 2020). The dynamics include a fluctuating line tension term added to Eq. (3):

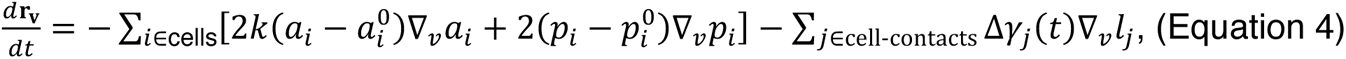

where *l*_j_ is the length of the cellular contact *j*, and the fluctuating line tension on contact edge *j* evolves according to Ornstein-Uhlenbeck scheme as

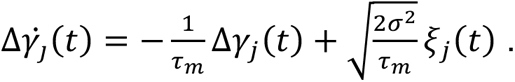

Here, the noise *ξ*(*t*) obeys W*ξ*_j_(*t*)X = 0 and W*ξ*_j_(*t*)*ξ*_*l*_(*t*^′^)X = *δ*_j*l*_*δ*(*t* − *t*^′^) with *τ*_*m*_ and *σ*^2^ are its relaxation time scale and long-time variance, respectively.

We melt each configuration for 3000 time units using *σ* = 0.5, which is followed by a sharp quench (*σ* = 0) and minimization over an additional 7000 time units. During these two steps, we allow T1 transitions. After that, all four-fold vertices are resolved into small edges of length 10^−3^ which is followed by another 1000 time units of minimization without T1 transitions using a smaller integration time step Δ*t* = 10^−4^.

##### Cell diminution and reinsertion

Each prepared configuration undergoes a simulation consisting of the following steps:

1. **Initial minimization:** The system is further minimized for *t*_eq_ = 300 time units.
2. **Cell deletion:** The preferred area of the mitotic cell is gradually reduced over *t*_sh_ = 100 time units to a final value 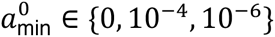 based on numerical experiments, using: 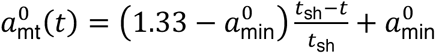. Subsequently, the preferred perimeter evolves as: 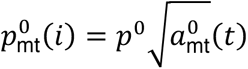, where *p*^0^ is the preferred perimeter of a normal cell.
3. **Post-deletion minimization:** The system is further minimized for 100 time units.
4. **Division and reinsertion:** The tiny cell is divided along an arbitrary axis passing through its centroid. The two daughter cells generated due to the cell division are grown together linearly over the duration of *t*_rg_ up to a size equal to half of that of the mitotic cell with their preferred areas evolving as: 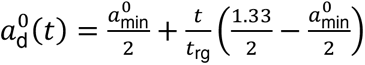 The time of division marks the reinsertion time. Here, the preferred perimeter evolves as: 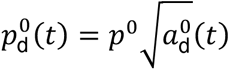.
5. **Final minimization:** The system is further minimized for an additional 300 time units.

##### Displacement profile calculation

Mitotic activity-induced displacement profiles are computed using the following steps:

We prepare one ordered 2D tissue (honeycomb lattice) by packing *N*_*c*_ = 1024 identical hexagonal cells, each with preferred area *a*^0^ = 1 and preferred perimeter *p*^0^ = 3.722, with periodic boundary conditions. The system is minimized for 300 time units, and the location of cell centroids (pre-deletion centroids) is recorded.

A single cell is arbitrarily chosen as the mitotic cell, which is then deleted using the same protocol described above with 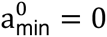. Following post-deletion minimization, new centroids are computed. The deletion-induced displacement map is obtained by subtracting the pre-deletion centroids from these newly computed centroids. Up to this step, we do not allow any T1 transition.

Next, we perform an additional 100 time units of minimization, allowing the T1 transition. This is followed by a division of the tiny cell such that the common cell contact between two daughter cells lies approximately parallel to the x-axis. Subsequently, the daughter cells are regrown, and then the system is minimized using the same “division and reinsertion” and “final minimization” protocol described earlier without T1 transformation. The final centroid locations are again recorded, and the reinsertion-induced displacement map is computed by subtracting centroid positions immediately after division from those after final minimization.

Original code underlying these simulations can be accessed from GitLab Repository at https://gitlab.com/tanmoy_jsi/basal_diminution

#### Intestinal organoid model

##### Mouse strains

The mice strains used in this study were a gift from Liheng Li’s lab. In all experiments involving membrane visualization with cell membrane-localized tdTomato (mT), mT/mG (JAX: 007676) (Muzumdar et al., 2007) was used. In experiments where the nuclei were visualized in addition to the cell membrane, mT/mG mice were crossed with R26-M2rtTA; TetOP-H2B-GFP (JAX: 016836) (Foudi et al., 2009). All mice experiments were approved by the Institutional Animal Care and Use Committee of the Stowers Institute for Medical Research (Protocol ID: 2017-0179) and the University of Nebraska Medical Center (Protocol ID: 22-078-11-EP).

##### Organoid culture

The organoid cultures were derived from murine duodenal crypt. The intestinal crypts were dissociated by treating duodenal tissue with Gentle Cell Dissociation Reagent (Stemcell Technologies) at room temperature and repeated pipetting in PBS. Crypts were enriched by passing the supernatant through a 70µm filter to exclude larger pieces of the intestinal tissue. The isolated crypts were seeded in Matrigel (Corning) and cultured in IntestiCult Organoid Growth Medium (Stemcell Technologies) at 37°C) and 5% CO2. The organoids were passaged every 7 days by dissociating and reseeding the intestinal crypts from the 3D culture. For tracking the nuclei in organoids, the expression of H2B-GFP in R26-M2rtTA; TetOP-H2B-GFP; mT/mG was induced by treating with Doxycycline (2 µg/ml) for 24 h.

##### Imaging organoid epithelium

Organoid epithelia were fixed with 4% PFA for 30 min and washed with PBST (1X PBS with 0.1% TritonX-100) before incubation with primary antibody was incubated overnight at four degrees. The secondary antibody was incubated for 2h at RT. The samples were mounted in SlowFade Diamond Antifade.

Following dilutions were used for staining the organoid tissue:

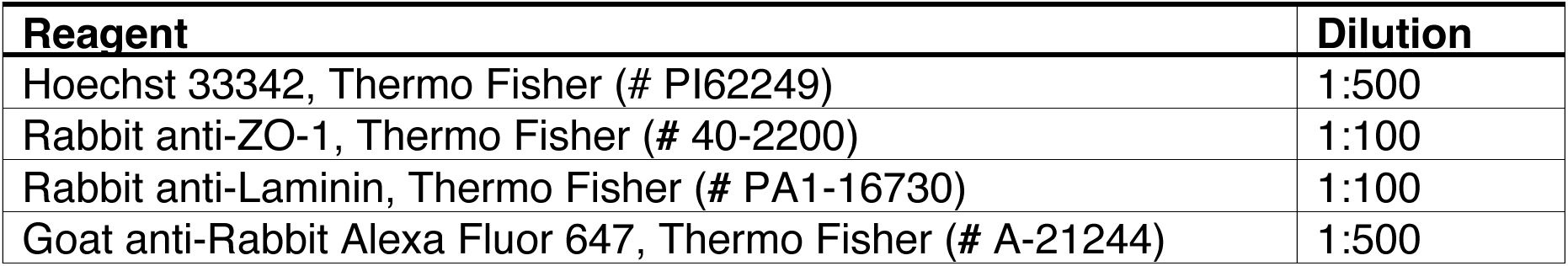

An LSM 900 Airyscan confocal microscope with Plan Apo 40x/1.4 Oil DIC (UV) V-IR and C-Apo 40x/1.2 W objectives was used for point scanning applications. Spinning disk image acquisition was performed using Nikon Ti Eclipse with Yokogawa CSU-W1 spinning disk head equipped with a Hamamatsu Flash 4.0 sCMOS and Plan Apo 40X 1.15 NA LWD. Additionally, the organoids were maintained at 37C and 5% CO2 during live imaging using stage-top incubation chambers from PeCon for Zeiss and OkoLab for Nikon. Timelapse images were acquired over time intervals of 4 mins and z sections of 1 µm. For imaging the basal regions of the organoid, the range of z started at the basal-most part of the organoid and ended at the apical-most region of the nucleus. In experiments involving mitotic arrest, organoids were treated with 5 µg/ml STC.

###### Image Processing

Adjustments to image contrast, brightness, and size were performed by linear interpolation and applied to the whole image using FIJI. Drift correction in time-lapse images was performed using the StackReg FIJI plugin. Images were compiled with Adobe Illustrator CC.

### Quantification and Statistical Analysis

#### Quantification

All quantifications including cell perimeter, area, cell-contact angles, and neighbor number were determined by manually segmenting cells using FIJI. Circularity was calculated using the following formula: *C = 4πA/P^2^*, where *A* and *P* are cell area and perimeter, respectively. Unless otherwise mentioned, images three timeframes prior to the initiation of basal diminution of mitotic cells were used for all quantifications relating to late interphase cell shape.

#### Statistical analyses

Sample sizes were based on the standard of the field. At least 20 mitotic events from at least five different organoids were tracked for each experimental condition. In the boxplots, circles represent mean values from individual cells, unless specified. The diamond box contains 25–75% percentiles of the data and the bar denotes the median. Wilcoxon Signed Ranks Test was used to compare matched samples (for example, same cell before and after mitosis) and Mann-Whitney U test was used to compare two independent samples. All statistical significance tests were carried out using Origin (OriginLab).

## CONTRIBUTIONS

S.P.R. designed the project, planned and performed experiments, analyzed data, provided mentorship, secured funding, and wrote the manuscript. T.S. planned and performed experiments, analyzed data, and contributed to manuscript editing. M.K. provided mentorship, acquired funding, and contributed to manuscript editing. R.M., A.C., and C.Z. conducted experiments and participated in data analysis. M.C.G. provided mentorship, secured funding, and contributed to manuscript editing.

## ACKNOWLEDGEMENTS

This work was supported by funding from multiple institutions and agencies. S.P.R. acknowledges support from a UNMC startup grant and NIH/NIGMS (P20 GM121316). T.S. and M.K. were supported by the Slovenian Research and Innovation Agency through research projects J1-3009 and J1-60013, the development funding pillar RSF-0106, and research core funding P1-0055. M.C.G. acknowledges support from the Stowers Institute for Medical Research.

---

**Figure S1.**
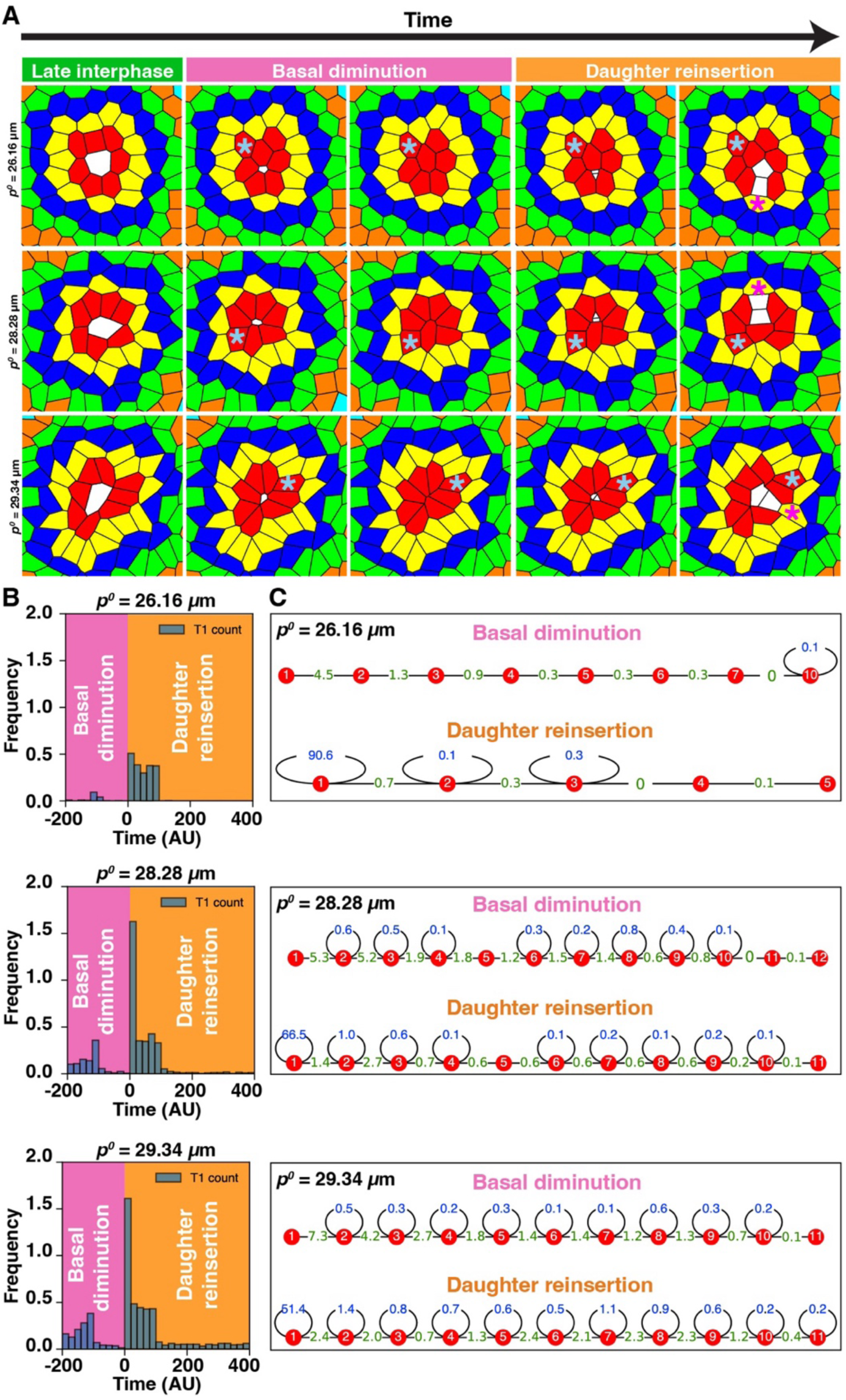
Vertex model simulations predict mitosis-induced basal cell-contact remodeling across a range of mechanical regimes. (A) Snapshots of cell divisions in tissues with varying 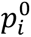 (26.16*μm*, 28.28*μm*, and 29.34*μm*). Mitotic cell: white; color-coded neighbors as in Fig. 1C. Asterisks mark T1 transitions. (B) Temporal distribution of T1 transitions for each 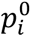. Negative and positive times indicate diminution and reinsertion, respectively (n = 324). (C) Spatial distribution of T1 transitions as a function of topological distance from the mitotic cell.

**Figure S2.**
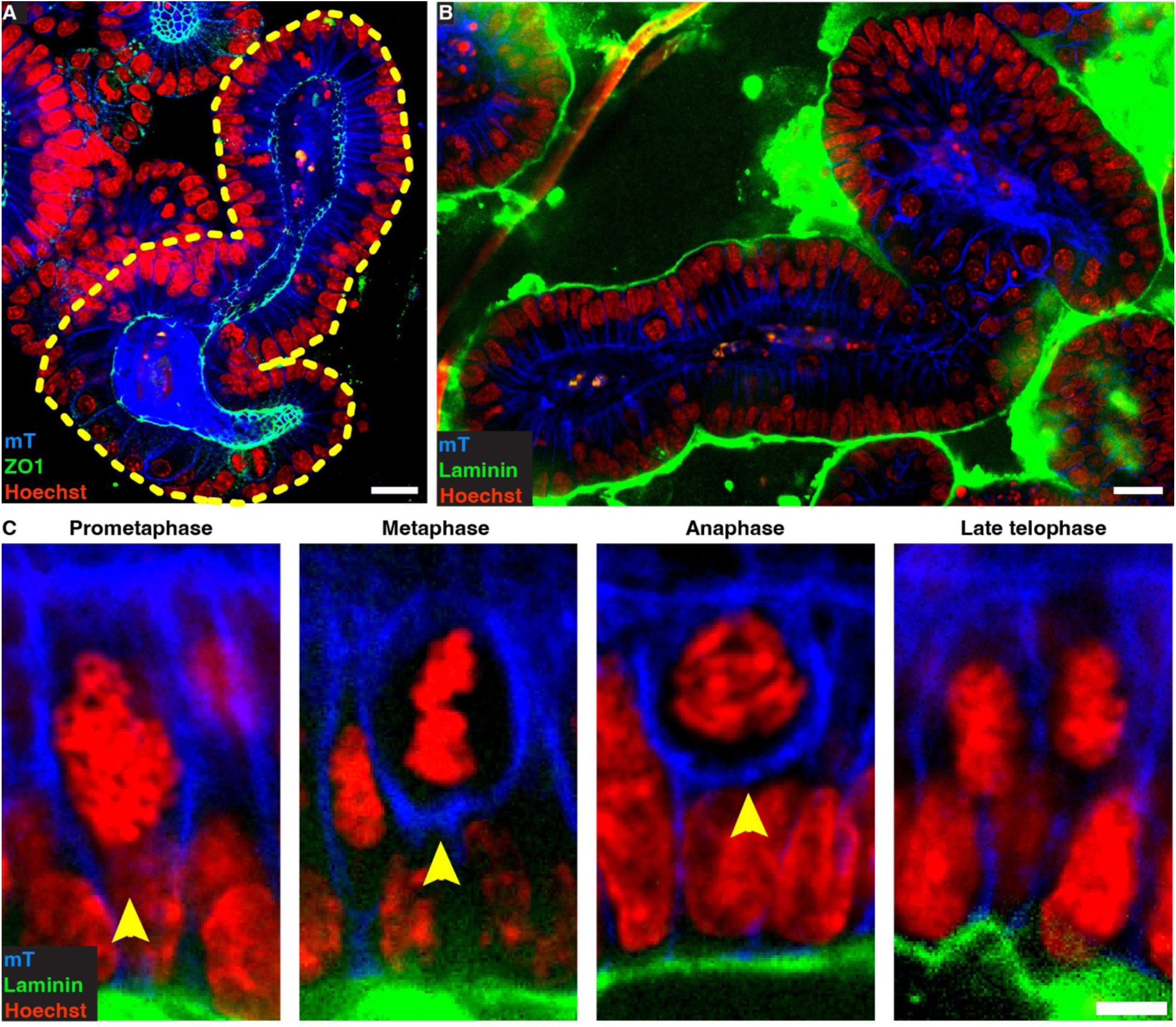
Proliferating epithelial cells from intestinal organoid culture are translocated from the basal region during the start of prophase and reinserted back into the basal region after cytokinesis. (A, B) Intestinal organoid epithelium strained for ZO1 (*n* = 16 organoids) to identify the apical domain (A) and laminin (*n* = 14 organoids) to identify the basement membrane (B). The yellow outline marks the organoid boundary. All scale bars, 20 µm (C) Mitotic cell body (arrowheads) is displaced away from the basement membrane at the start of mitosis. After cytokinesis, the daughter cell bodies return to the basal plane.

**Figure S3.**
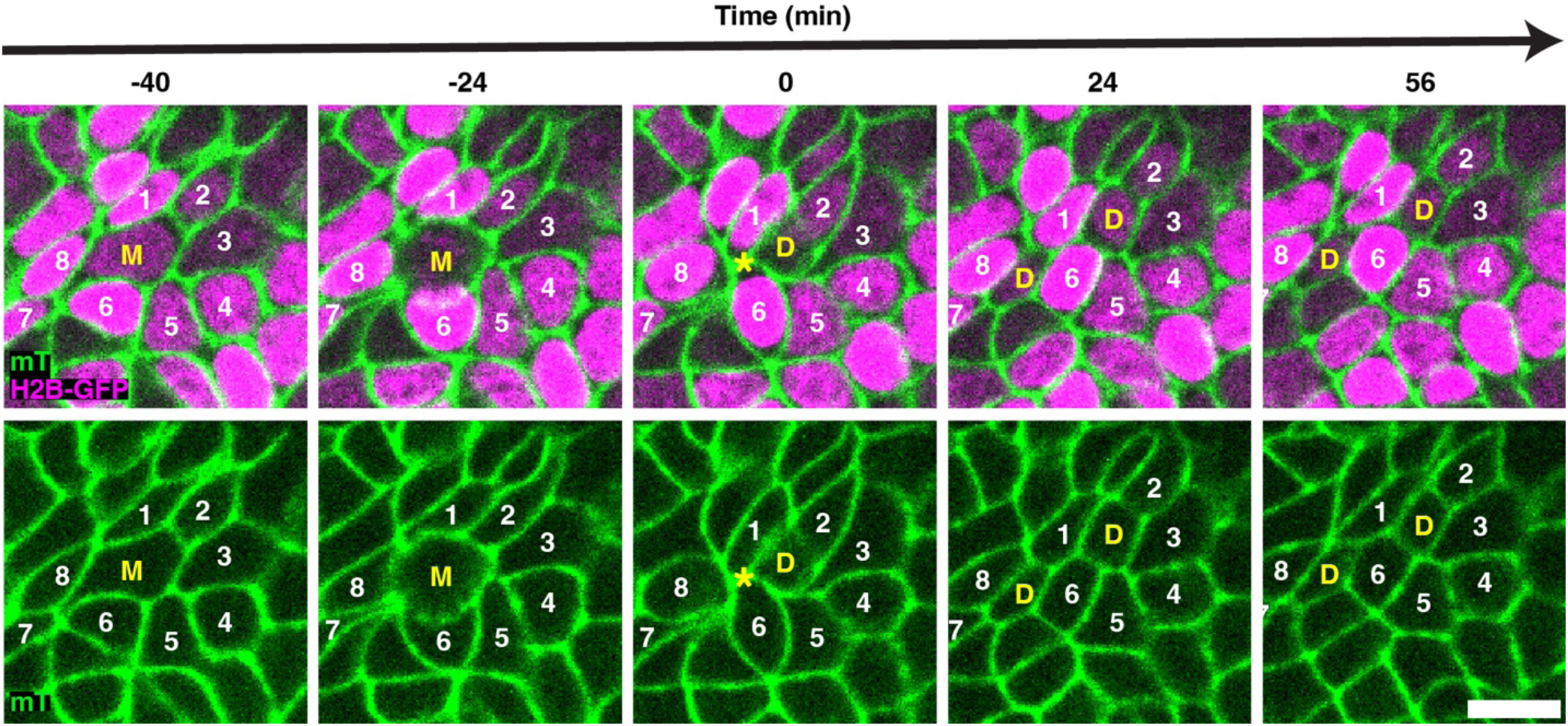
Daughter cell separation at the basal plane is an infrequent topological outcome. Timelapse images showing the basal cell-contact topology of a mitotic cell (‘**M’**), its immediate neighbors (‘**1-8’**), and daughter cells (‘**D’**). Membranes: mTomato (green); nuclei: H2B-GFP (magenta). Scale bar, 10 µm.

**Figure S4.**
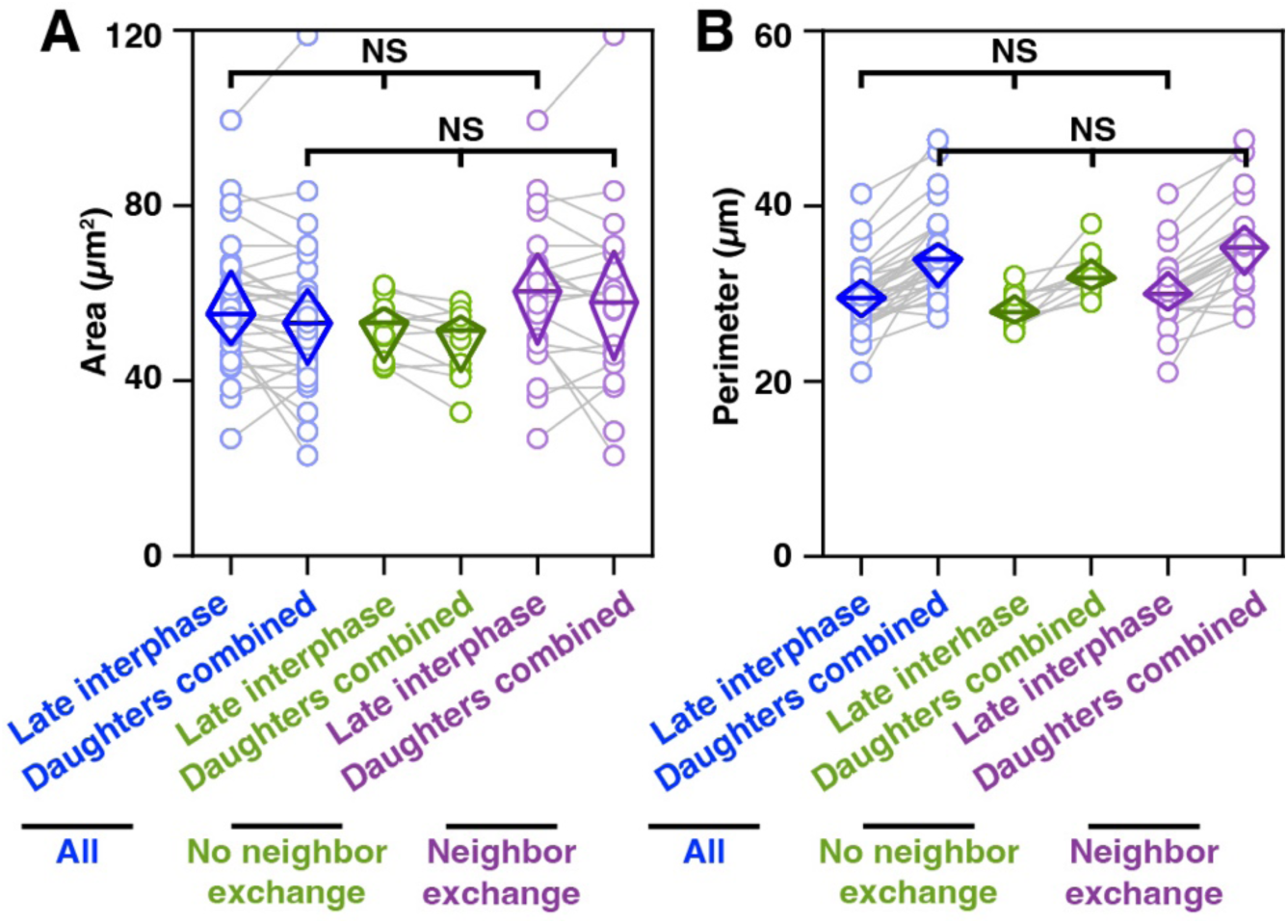
Basal surface area and perimeter of mitotic cells remain conserved across neighbor exchange conditions. Mother cell area (A) and perimeter (B) before mitosis compared to that of the daughter cells combined. The data is segregated based on whether (green) or not (magenta) neighbors exchanged after daughter cell reinsertion into the basal region. Daughters from the same mother are paired. *n* = 31 untreated and 23 STC-treated mitotic cells. NS denotes *p*-value > 0.05, Wilcoxon Paired Signed Ranks Test.

**Figure S5.**
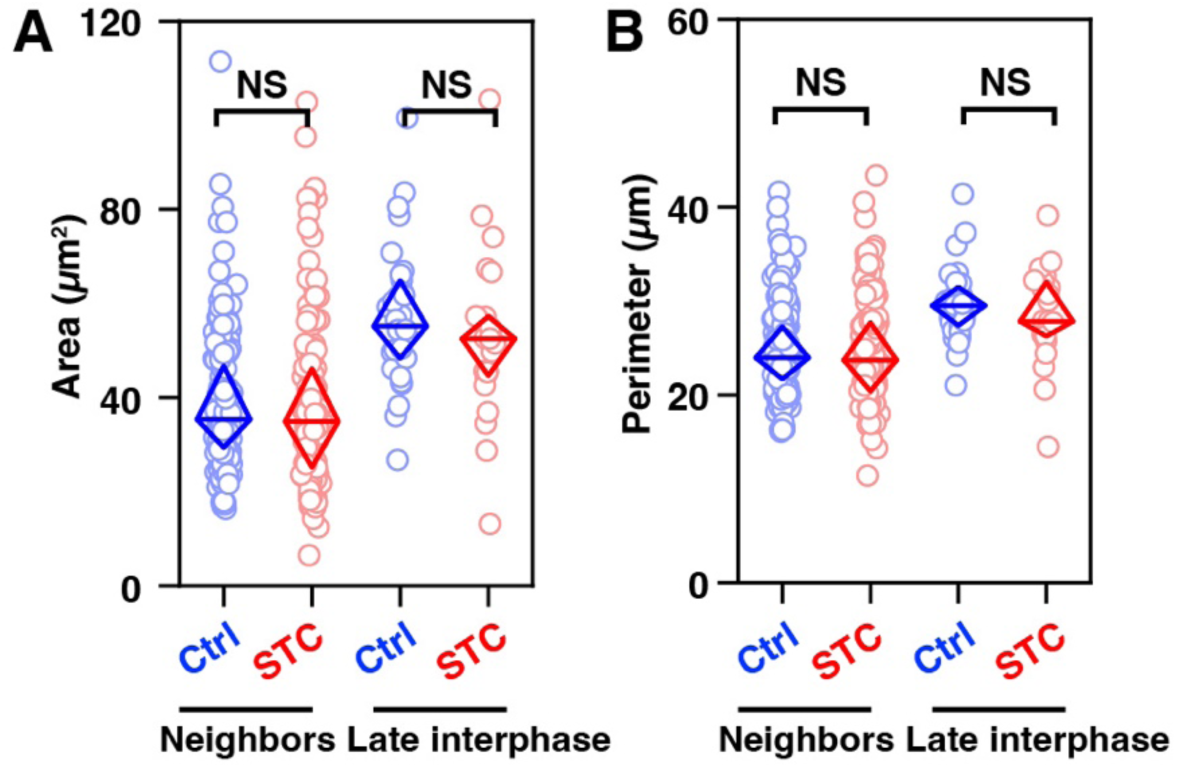
The morphology of STC-treated interphase cells is unaltered. Basal area (A) and perimeter (B) of neighbors of untreated (blue) and STC-treated (red) late interphase cells and their immediate neighbors. *n* = 31 untreated cells and 23 STC-treated prophase cells. NS denotes *p*-value > 0.05, Mann-Whitney Test.

